# Identifying cellular RNA-binding proteins during infection uncovers a role for MKRN2 in influenza mRNA trafficking

**DOI:** 10.1101/2023.10.30.564690

**Authors:** Stefano Bonazza, Hannah Coutts, Swathi Sukumar, Hannah Turkington, David G Courtney

## Abstract

Utilisation of RNA-binding proteins (RBPs) is an important aspect of post-transcriptional regulation of viral RNA. Viruses such as influenza A viruses (IAV) interact with RBPs to regulate processes including splicing, nuclear export and trafficking, while also encoding RBPs within their genomes, such as NP and NS1. But with almost 1000 RBPs encoded within the human genome it is still unclear what role, if any, many of these proteins play during viral replication. Using the RNA interactome capture (RIC) technique, we isolated RBPs from IAV infected cells to unravel the RBPome of mRNAs from IAV infected human cells. This led to the identification of one particular RBP, MKRN2, that associates with and positively regulates IAV mRNA. Through further validation, we determined that MKRN2 is involved in the nuclear-cytoplasmic trafficking of IAV mRNA likely through an association with the RNA export mediator GLE1. In the absence of MKRN2, IAV mRNAs accumulate in the nucleus of infected cells, which we suspect leads to their degradation by the nuclear RNA exosome complex. MKRN2, therefore, appears to be required for the efficient nuclear export of IAV mRNAs in human cells.

## Introduction

In recent years, our understanding of the RNA-binding protein (RBP) interactome has become significantly more informed. The development of methods such as RNA-interactome capture (RIC), enhanced RIC (eRIC) and RNA affinity pulldown mass spectrometry (RAP-MS), has allowed researchers to further elucidate the true RBPome of total mRNAs, as well as individual RNAs, in a given cell population.^1–4^ These techniques, in addition to viral cross-linking and solid-phase purification (VIR-CLASP),^5^ have now been used to good effect to identify the RBPome of viral RNAs from Sindbis virus,^6,7^ Chikungunya virus (CHIKV),^5^ and, in multiple studies, SARS-CoV-2.^8–10^

It has been reported previously, in cell culture, that influenza A virus (IAV) mRNAs can comprise up to 50% of the total mRNA population of an infected cell.^11^ Therefore, we reasoned that use of the RIC method, resulting in the purification of total mRNAs and their interacting proteins, would be ideal for the identification of pro-viral host RBPs potentially hijacked during IAV replication. Unravelling the IAV RBPome is essential to better understand the post-transcriptional regulation of IAV mRNAs. Additionally, this knowledge is critical for a better comprehension of the properties surrounding viral RNA stability, nuclear-cytoplasmic trafficking, and translation, all of which are governed by RBPs.^12^ In particular, this method could provide a more thorough understanding into how intronless IAV mRNAs escape the nucleus after transcription. Though this is still an open question, a recent study by Bhat et al (2023) has shed some light on this process of nuclear-cytoplasmic trafficking. The authors reported that the TREX-2 complex is recruited to viral mRNAs, then through an association with NXF1, the viral mRNAs are efficiently exported via the nuclear pore complex (NPC) to the cytoplasm.

Herein, we aimed to unravel novel host factors important in the post-transcriptional processing of IAV mRNA. Indeed, we identified numerous RBPs that, when their expression was manipulated, had a negative impact on viral replication kinetics. This work concentrates primarily on the understudied MKRN2 protein, a member of a family of Makorin zinc finger domain-containing proteins, that encodes for 4 classical C_3_H zinc finger RNA binding domains, an RNA binding Makorin Cys-His domain, and a RING-type zinc finger domain.^13^ MKRN2 has previously been implicated in mouse infertility, and cancer cell proliferation, while one recent report revealed its involvement in mRNA nuclear trafficking in zebrafish.^14–19^

In this study we uncover differences in the RBPome of poly(A)+ RNA upon infection with different strains of H1N1 IAV. We reveal the importance of one of these RBPs, MKRN2, in the post-transcriptional processing of IAV mRNAs, specifically in regard to viral mRNA nuclear export. We demonstrate that GLE1, a proven interactor of MKRN2 and a component of mRNA nuclear export machinery elicits the same effects on viral mRNAs. Finally, we speculate that MKRN2 may act as an mRNA export adapter protein to recruit factors such as FYTTD1 and the nuclear export complex, leading to the efficient export of viral mRNA.

## Results

### RIC methodology effectively identifies RBPs in influenza A virus-infected cells

To initially determine the RBPome of mRNAs from influenza A virus-infected cells, we employed the RIC approach referred to previously. This technique allows for the purification of all proteins bound to poly(A)+ RNA through conventional UV crosslinking, followed by mass spectrometry (Fig 1A). Minimally, influenza A virus encodes for 10 distinct poly(A)+ mRNA (Fig 1B). Following pulldown of all poly(A)+ mRNA and covalently crosslinked proteins, from mock, A/WSN/33 infected or A/California/7/2009 infected A549 cells, a silver stain was performed to confirm the presence of a diverse protein population suitable for mass spectrometric analysis (Fig 1C). Indeed, we suspected that viral NP protein was visible as a band at approximately 55kDa on the A/WSN/33 pulldown sample, and to a lesser extent in the A/California/7/2009 pulldown sample. Additionally, Western blot analysis confirmed the viral RBP NP was present only in infected samples, as well as in pulldown samples (Fig 1D). The presence of a known cellular mRNA RBP, YTHDF2, was confirmed in all RIC samples, while ACTB, not believed to be an RBP, was also blotted as a RIC negative control (Fig 1D). Finally, we confirmed by qPCR that the RIC samples contained substantial amounts of viral mRNAs, namely NP and NA mRNA, when compared to the mock, while the GAPDH and ACTB mRNAs were equally distributed among all 3 conditions (Fig 1E).

**Figure 1 –.**
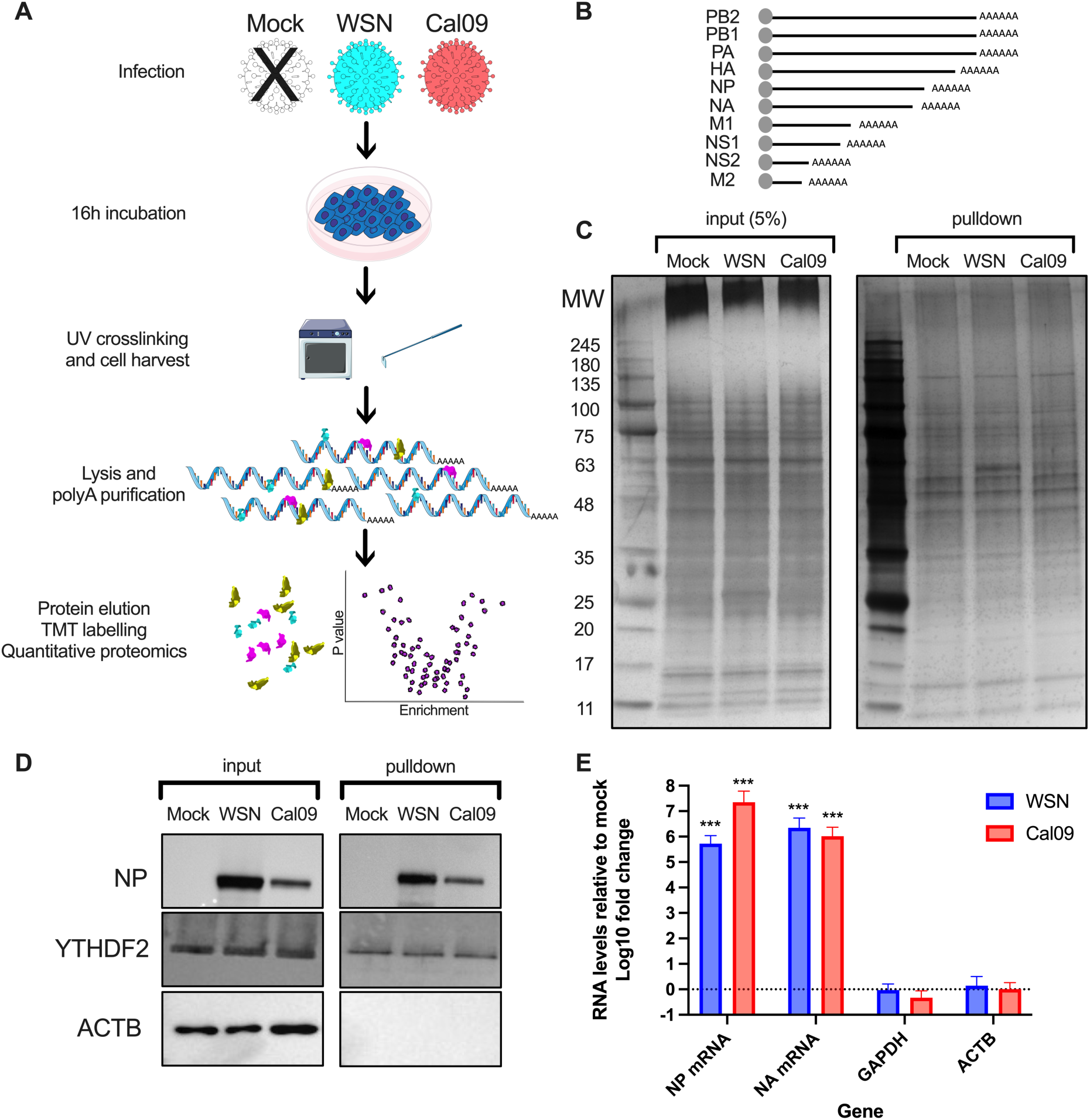
Collection of RIC samples for the identification of IAV mRNA specific RNA-binding proteins. (A) Schematic of the workflow for RIC on IAV infected A549 cells. (B) Representation of the minimal species of A/WSN/33 mRNAs expressed during infection. (C) Silver stain of the input and RIC pulldowns, showing the distribution of proteins present in the RIC samples and, likely, the viral NP protein appearing as a band of 55kDa in the A/WSN/33 sample, and to a lesser extent in the A/California/7/2009 sample. (D) Western blot validation of the RIC pulldown, demonstrating, as expected, the presence of viral NP in A/WSN/33 and A/California/7/2009 samples only, a known RBP YTHDF2 found in all 3 samples, and ACTB not present in any pulldowns. (E) qPCR validation of NP and NA mRNAs highly enriched in RIC pulldowns, while GAPDH and ACTB are at comparable levels to Mock.

### Identification of the mRNA RBPome of influenza A virus infected cells

Upon TMT labelling, mass spectrometry and quantitative analysis of the 9 RIC samples on MaxQuant and Perseus, we determined the RBPome for each test condition. Additionally, we quantified enrichment of RBP abundance in A/WSN/33 over mock (Fig 2A), and A/California/7/2009 over mock (Fig 2B). Adjusted P values were also calculated, with a cutoff of p<0.05. NP was found to have the strongest P value of any RBP for both viruses, which is not surprising considering its strong RNA-binding capabilities. The same is true for NS1, which was the most highly enriched protein in the A/WSN/33 infected samples. For A/WSN/33 infected cells, the 3 influenza polymerase proteins, PB2, PB1 and PA, as well as HA and M1 were also found to be enriched, likely as these proteins are not present in the mock sample. Statistically, they could be misinterpreted as overrepresented in the A/WSN/33 RBPome. So, their inclusion here does not confirm or discount their potential to be mRNA binding proteins.

**Figure 2 –.**
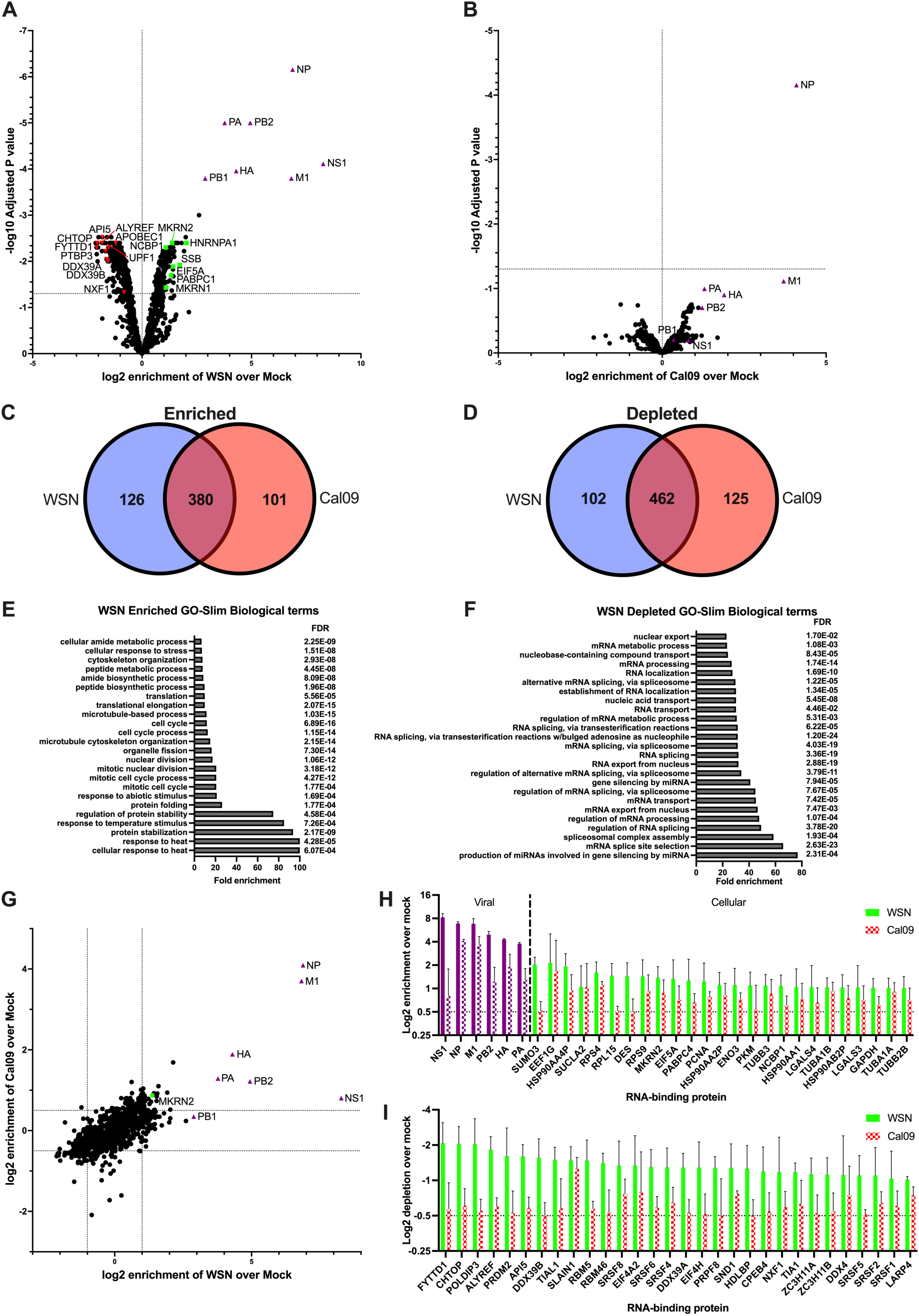
Mass spectrometry analysis of IAV mRNA-binding proteins and identification of enriched and depleted RBPs during infection. Volcano plots showing the enrichment profile of proteins isolated from RIC pulldowns from A/WSN/33 infected (A) and A/California/7/2009 infected (B) cells when compared to Mock. Venn diagrams displaying the proteins enriched or depleted on poly(A)+ RNA after infection with A/WSN/33 or A/California/7/2009, highlighting the commonly enriched proteins (C) and depleted proteins (D). Slim GO biological terms found to be enriched among the aforementioned enriched (E) and depleted (F) proteins from the RIC pulldown of A/WSN/33 infected cells over Mock. (G) A graph of all the proteins identified in both A/WSN/33 and A/California/7/2009 RIC eluates, plotted as log_2_ enrichment over Mock. MKRN2 enrichment is highlighted in green. Enrichment profiles of all proteins found to be greater than 0.5 fold enriched (H) or less than −0.5 fold (I) in both A/WSN/33 and A/California/7/2009 RIC samples, average between 3 biological replicates. Viral proteins have been coloured purple.

Altogether, we observed good concordance between the RBPs in the A/WSN/33 and A/California/7/2009 infected samples, with >75% and >78% of all enriched and depleted RBPs, respectively, found in both datasets. We then sought to better understand whether RBPs involved in specific pathways or processes are being enriched or depleted in these IAV infected samples. We generated a list of proteins that were altered by >0.5 fold in A/WSN/33 infected cells and determined the biological Slim GO term associated with these datasets. We found that the RBP dataset we identified as being enriched on poly(A)+ RNA in A/WSN/33 infected cells is predominantly comprised of proteins involved in cellular stress responses, such as temperature. In contrast, the dataset from depleted RBPs was comprised of proteins involved in splicing and mRNA transport. The reduction in the binding of RBPs related to splicing function in A/WSN/33 infected cells can be explained by the fact that only M and NS segment are spliced in A/WSN/33.

### Validation of potential pro-viral IAV mRNA binding proteins

To validate selected enriched RBPs from our RBPome dataset, we used a A/WSN/33 reporter virus expressing a PA-mNeon fusion protein in place of wild-type PA. Infected cells were identified by the presence of green fluorescence. siRNA mediated knockdown was performed in A549 cells for 8 different RBPs identified from our dataset, alongside a non-specific control siRNA (NSC), and a control siRNA targeting ALYREF, a cellular RBP involved in the export of spliced host mRNAs. An siRNA targeting FYTTD1, a known nuclear mRNA interacting protein, was also included. After 48h of siRNA knockdown, cells were infected with the reporter virus for 16h, before the visualisation of cells by fluorescence microscopy and protein extraction and Western blotting. Western blotting revealed a clear decrease in viral protein expression in cells depleted for SUMO3, HNRNPA1, MKRN2 (but not MKRN1) and EIF5A (Fig 3A). For the most part, this was consistent with the quantification of mNeon positive cells by fluorescence microscopy, with all 4 of these knockdowns almost completely abrogating WSN PA-mNeon protein expression (Fig 3B & C).

**Figure 3 –.**
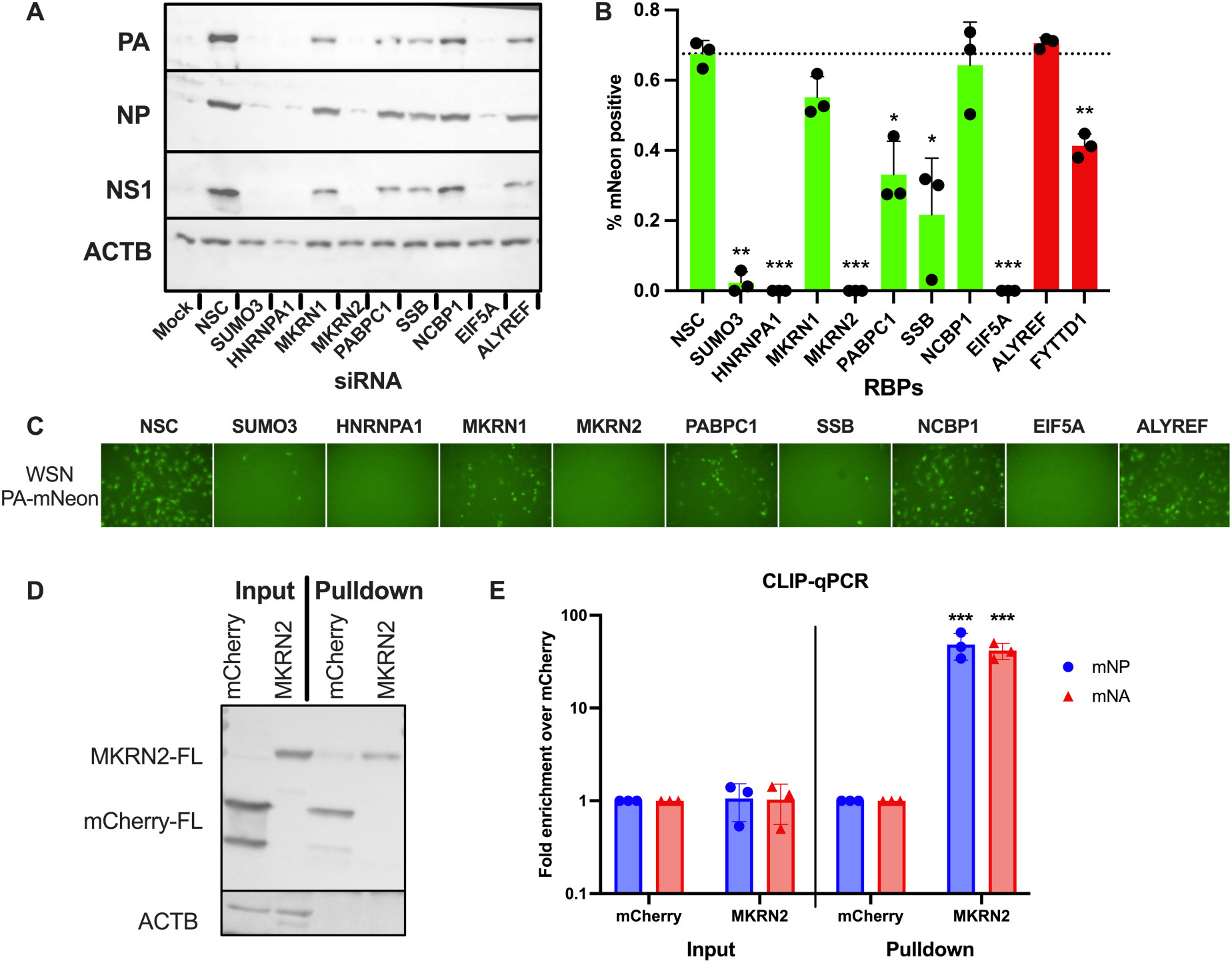
Validation of RBP hits and the confirmation of MKRN2 as an IAV mRNA-binding protein. A reporter A/WSN/33 virus expressing PA-mNeon was used to screen a number of RBPs for their proviral function. Western blot was performed to assess A/WSN/33 protein expression (A), while the percentage of green fluorescent cells was quantified by fluorescence microscopy (B). Representative images of infected siRNA treated cells are also shown (C). Further validation of the RNA-binding capabilities of one RBP, MKRN2, was performed. CLIP-qPCR of mCherry or MKRN2 in IAV infected cells was carried out. Input and pulldown samples were assessed by Western blot (D) while RNA isolated from the pulldowns was quantified by qPCR, where MKRN2 was found to readily bind NP and NA mRNAs.

Interestingly, the enriched RBPome dataset contained MKRN1 and MKRN2, two RBPs from the same protein family with similar structures. However, depletion of these proteins yielded very different effects on A/WSN/33 infection. We were curious as to the cause of this phenotypic difference, so we chose to proceed with the characterisation of MKRN2 and its role in IAV replication. Viral RNA-binding capabilities of MKRN2 were validated through CLIP-qPCR (Fig 3E). HEK293T cells expressing a C-terminally Flag-tagged mCherry or MKRN2 were infected with wild-type A/WSN/33 virus for 16h. Cells were then UV crosslinked, lysed and an immunoprecipitation was performed for Flag. Input and pulldown protein samples were assessed by Western blotting, demonstrating a purification of Flag-tagged mCherry or MKRN2 and the absence of ACTB in the pulldown (Fig 3D). Additionally, RNA was isolated from the input and pulldown samples, followed by qPCR to quantify the levels of NP and NA mRNA. Normalising to the mCherry input levels, we observed that the NP and NA mRNA input levels were comparable, while in the pulldown samples the viral transcripts were enriched by up to 60-fold (Fig 3E). These findings provided strong evidence for the RNA-binding capabilities of MKRN2 to IAV mRNAs in this context.

### MKRN2 as a novel pro-viral host factor

We sought to further our understanding of the pro-viral nature of MKRN2, and whether the observed relationship is conserved among additional IAV strains. After validation of the MKRN2 siRNA pool by Western blotting (Fig 4A), we performed multicycle infections of A549 MKRN2 depleted cells with A/WSN/33, the 2009 pandemic H1N1 A/California/7/2009 virus and an H3N2 A/Norway/466/2014 strain. MKRN2 knockdown resulted in a significant and reproducible reduction in viral replication of all three strains at 72 hpi (Fig 4B-D). As the A/California/7/2009 strain is more clinically relevant and similar to a bona fide and circulating human H1N1 seasonal strain of IAV, we decided to proceed with further characterisation of the mechanism of action of MKRN2 using the A/California/7/2009 virus. (Fig 4B-D).

**Figure 4 –.**
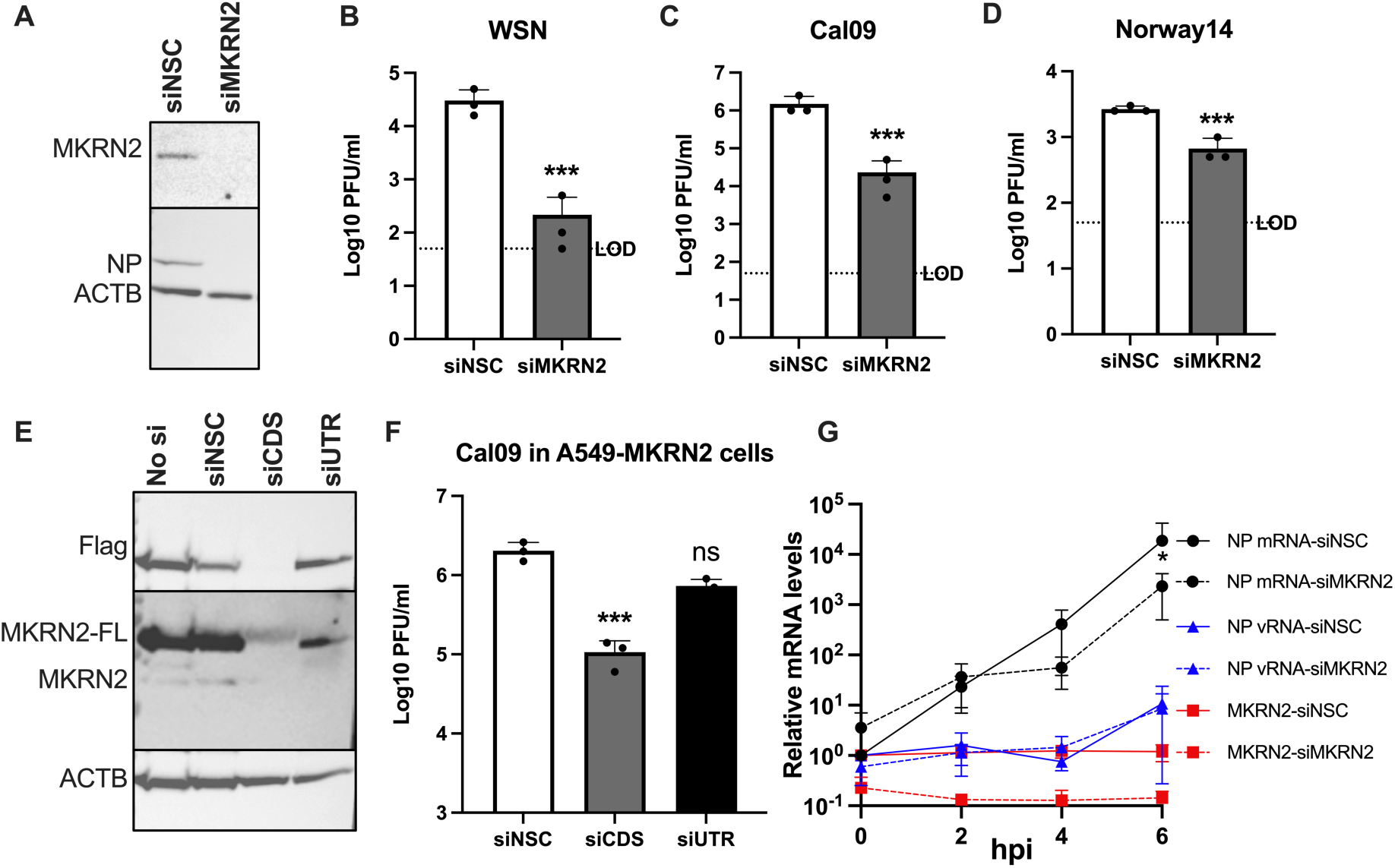
Investigation of the phenotypic effect of MKRN2 depletion on IAV replication. (A) Demonstration by Western blotting that the MKRN2 siRNA pair reduce MKRN2 protein levels at 48 h post transfection. Knockdown of MKRN2 in A549 cells, followed by infection with A/WSN/33 (B), A/California/7/2009 (C) or A/Norway/466/2014 (D) significantly reduced virus output from these cells as measured by plaque assay. (E) Western blot demonstrating that the MKRN2 siRNA pair reduce expression of endogenous and lentiviral expressed MKRN2 in an overexpression A549 cell line, while an siRNA pair targeting the MKRN2 3’ UTR only reduce expression of the endogenous MKRN2 protein. (F) A/California/7/2009 titres from MKRN2 overexpression A549 cells demonstrate that while reducing all MKRN2 expression significantly reduced virus output, knocking down endogenous MKRN2 while expressing wild-type MKRN2 from a lentiviral cassette rescues this phenotype. (G) Early viral RNA dynamics in siMKRN2 treated cells were quantified by qPCR, revealing that a loss of MKRN2 significantly reduced viral NP mRNA levels early in infection.

To confirm if the observed phenotypes are MKRN2 specific, we generated a stably expressing Flag-tagged MKRN2 overexpression A549 cell line via lentiviral transduction and single cell cloning. We aimed to use this cell line as a rescue system. Western blotting confirmed that the Flag-tagged and endogenous wild-type MKRN2 were expressed in untransfected and siNSC transfected cells (Fig 4E). The MKRN2 siRNA used throughout, termed here as siCDS due to its targeting of the coding sequence, successfully knocked down the expression of both forms of MKRN2 (Fig 4E). Whereas a newly designed siRNA targeting the 3’ UTR of MKRN2, termed siUTR, that is only present on the wild-type mRNA, depleted only the endogenously encoded MKRN2 (Fig 4E). The increased levels of MKRN2 did not alter the replication dynamics of A/California/7/2009 (data not shown). Similarly, to Fig 4C, we observed a robust decrease in viral titres from MKRN2 overexpression cells when overall MKRN2 protein levels are reduced. In contrast, when only the endogenous wild-type MKRN2 protein is reduced, we see no significant reduction in viral titre, likely due to the ongoing expression of Flag-tagged MKRN2 (Fig 4F). This confirmed that the siRNA mediated reduction in viral titres observed was MKRN2 specific.

### MKRN2 expression is positively correlated with IAV mRNA levels

Given the global reduction in IAV gene expression and replication in MKRN2 depleted cells, we proceeded to investigate which stage of the viral replication cycle is likely affected by the loss of MKRN2. To accomplish this, we performed single cycle growth curves and assessed A/California/7/2009 NP mRNA and vRNA levels by qPCR. We detected a significant reduction in NP mRNA in siMKRN2 treated cells at 6 hpi, however, no difference in NP-vRNA levels was observed at any time point (Fig 4G). This phenotype was recapitulated in A/WSN/33 growth curves, where viral mRNA levels in a single round infection were significantly lower when MKRN2 was depleted (Supplementary Fig 2). This suggests a role of MKRN2 in regulating viral mRNA early in infection.

### MKRN2 nuclear relocalisation upon infection affects IAV mRNA levels

As we observed this inhibitory phenotype early in infection, we wanted to better understand the role MKRN2 may play at this stage of the IAV replication cycle. We therefore decided to visualise the subcellular distribution of MKRN2 over the course of an infection. The localisation of MKRN2 and NP were imaged at 0, 4 and 8 hpi (Fig 5A). It was initially observed that MKRN2 has a uniform distribution in A549 cells across both the cytoplasm and the nucleus, amounting to an average nuclear fraction of 0.24 (Fig 5B), and remaining mostly unchanged at 4 hpi. As expected, the percentage of NP protein in the nucleus increased during the course of the infection (Fig 5C). However, at 8 hpi, we observed a significant shift in MKRN2 distribution from the cytoplasm to the nucleus amounting to on average an ∼0.35 nuclear fraction of MKRN2 protein This constitutes an almost 50% increase in the nuclear fraction of MKRN2 in infected cells, and hints at a role for MKRN2 in the processing of IAV mRNA in the nucleus.

**Figure 5 –.**
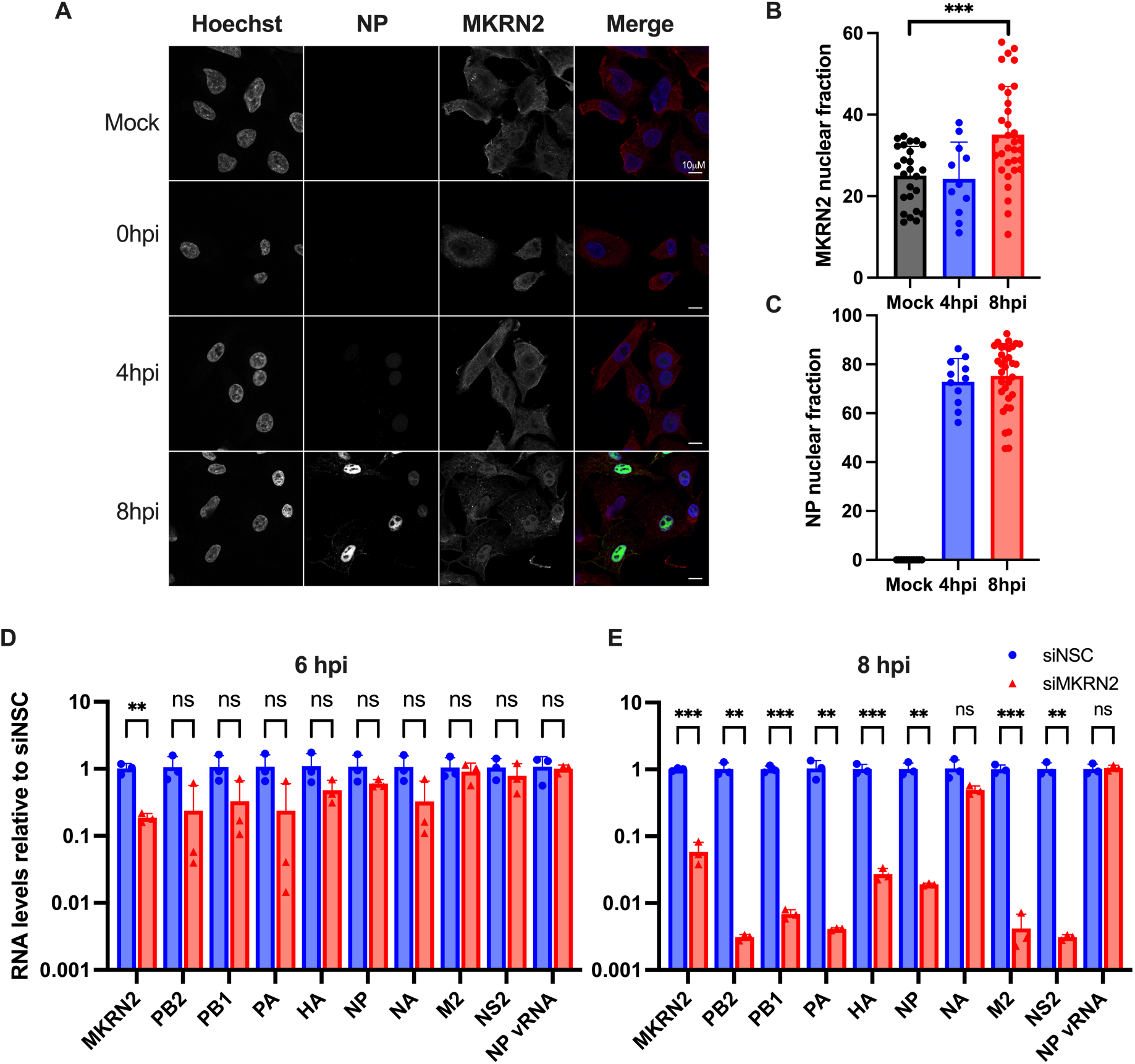
MKRN2 subcellular localisation during IAV infection and its role on viral mRNA dynamics. (A) Immunofluorescence staining and imaging was performed over a A/WSN/33 infection time course to visualise MKRN2 localisation. (B) The subcellular localisation of MKRN2 was quantified from these images and the nuclear fraction of all MKRN2 was calculated. (C) The NP nuclear fraction was quantified in a similar manner. The levels of primary A/California/7/2009 transcripts in infected control and MKRN2 depleted cells were quantified by qPCR at 6 hpi (D) and 8 hpi (E) alongside NP vRNA and MKRN2 mRNA levels.

With this clear relocalisation of MKRN2 occurring after 4 hpi, we analysed the effect of MKRN2 depletion on primary influenza mRNA transcripts after this timepoint. For this, we halted translation via cycloheximide treatment at 1 hpi and quantified the levels of 8 A/California/7/2009 mRNAs by qPCR at 6 and 8 hpi (M1 and NS1 cannot be quantified without also detecting M2 and NS2 transcripts, respectively). As expected, MKRN2 mRNA levels were significantly depleted in siMKRN2-treated cells, while NP vRNA levels were unaltered at both timepoints (Fig 5D). At 6 hpi we did not observe a significant change in the levels of any IAV transcripts, though they were all on average decreased in siMKRN2 treated cells compared to siNSC treated cells (Fig 5D). However, by 8 hpi, the primary IAV transcripts in the MKRN2 depleted cells were significantly decreased when compared to NSC treated cells, except for NA mRNA, which on average was lower but not significantly reduced (p=0.06) (Fig 5E). Between 6 and 8 hpi, the levels of IAV transcripts were not significantly altered in the control, as we believe primary transcription is likely to have peaked (Supplementary Fig 3). Due to the differences in IAV mRNA levels between 6 and 8 hpi in depleted cells, we hypothesise that IAV mRNAs may be accumulating in the nucleus of MKRN2 treated cells, likely leading to the degradation of viral mRNAs by the nuclear RNA exosome machinery. Therefore, we sought to uncover how the presence of MKRN2 may prevent the degradation of viral mRNAs.

### Nuclear export of IAV mRNA is inhibited by MKRN2 depletion

One recent study by Wolf et al. (2020), performed immunoprecipitation-mass spectrometry (IP-MS) of GFP tagged human RNA-binding E3 ubiquitin ligases, including MKRN1, MKRN2 and MKRN3. We acquired and reanalysed the raw MS data to compare protein interactors between these three MKRN-family RBPs. Given the discrepancy between MKRN1 and MKRN2 in reducing IAV expression (Fig 3A & B), we were interested in determining the specific interactome of each protein. In total, all 3 MKRN proteins were found to interact with a common set of 226 proteins. MKRN1 and MKRN3 interacted with 3 unique proteins, while MKRN2 had 7 unique interactors (Fig 6A), 5 of which were in common with the original published analysis. STRING analysis of the latter set of proteins uncovered that the 7 interactors complex together (Fig 6B) and are generally found to be involved in RNA transport (Fig 6C). This finding, in conjunction with the observations of Wolf et al. (2020) highlighted that MKRN2 association likely affects mRNA nuclear export. Alongside the relocalisation of MKRN2 during infection this prompted us to investigate if MKRN2 aids IAV mRNA export, which would in turn prevent their degradation mediated by nuclear retention.

**Figure 6 –.**
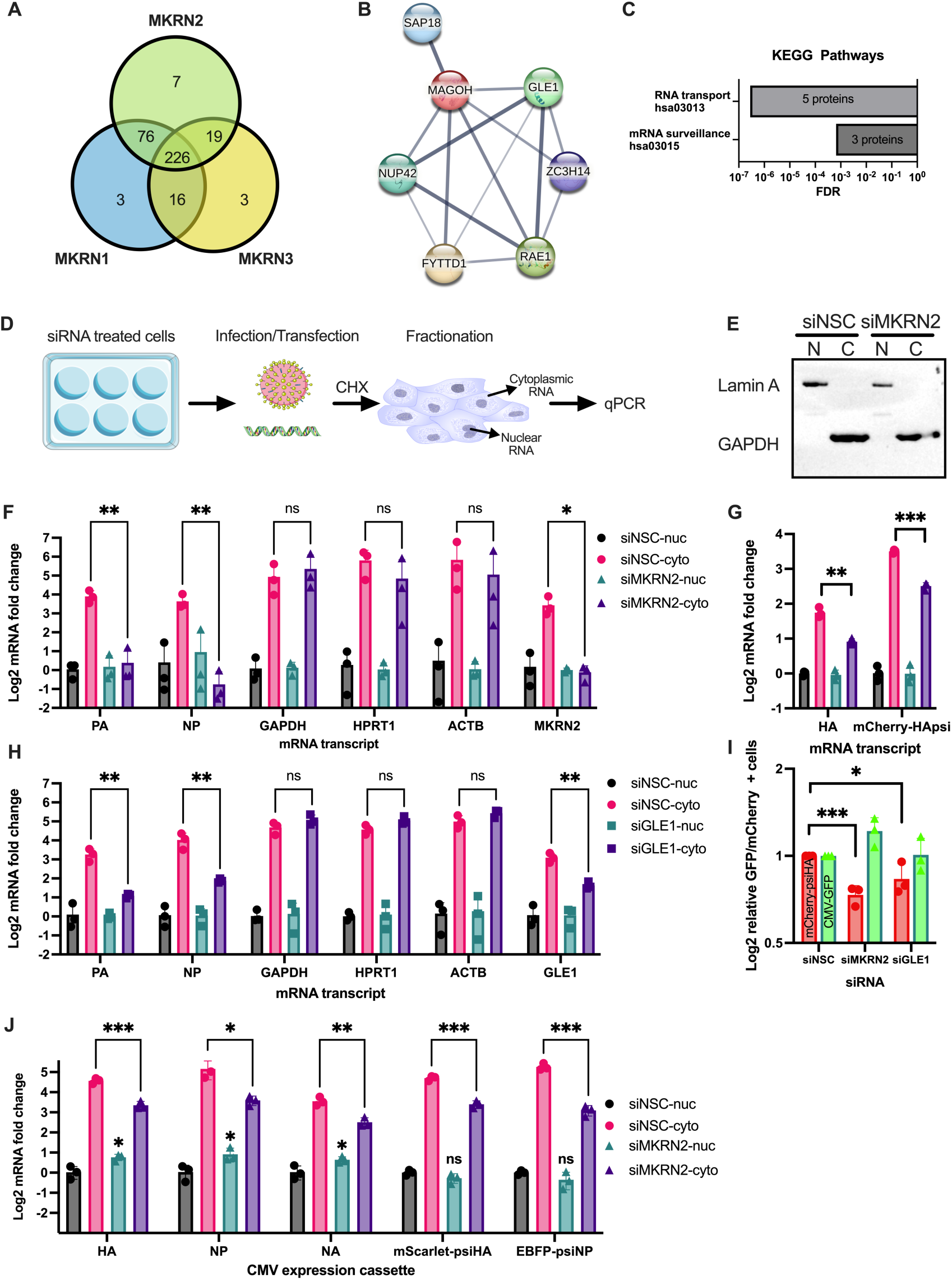
MKRN2 is involved in the nuclear export of viral mRNAs in association with GLE1. (A) Our reanalysis of IP-MS data from Wolf et al (2020) uncovered a large degree of overlap between MKRN-interacting proteins as well as some unique interactors. For MKRN2 the 7 unique interactors were found to cluster together by STRING (B) and a KEGG analysis revealed that these proteins were predominantly involved in RNA transport (C). (D) Schematic of siRNA treatment, infection and fractionation to uncover the nuclear and cytoplasmic fractions of IAV and cellular mRNAs. (E) Representative Western blot validation of the fractionation efficiency of our protocol. (F) By qPCR the levels of viral and cellular mRNAs in nuclear and cytoplasmic fractions were quantified and normalised to the control nuclear fraction. (G) A mini-replicon assay demonstrated that inclusion of the HA packaging signal was sufficient for MKRN2 depletion to down regulate the nuclear export of the mCherry pseudo segment mRNA. (H) Nucleo-cytoplasmic fractionation was performed on GLE1 depleted cells, similar to F where PA and NP viral transcripts were found to have depleted cytoplasmic fractions. (I) A mini-replicon mCherry reporter assay was performed in siRNA treated cells, where the number of mCherry and GFP positive cells were measured. Data was normalised to the siNSC treated cell numbers. (J) Nucleo-cytoplasmic fractionation was again performed, in this instance on 293T cells transfected to express transgenes from a CMV promoter all with IAV packaging signals. MKRN2 depletion was found to still inhibit cytoplasmic levels, even in this polII expression system.

To this end, we performed a nucleo-cytoplasmic fractionation followed by qPCR in siRNA transfected cells infected with A/California/7/2009 for 8h and treated with cycloheximide at 1 hpi (Fig 6D). The purity of the nuclear and cytoplasmic fractions was confirmed by Western blotting for Lamin A/C (nuclear) and GAPDH (cytoplasmic) (Fig 6E). qPCR analysis revealed an approximately 4-fold increase in the concentration of PA and NP mRNA in the cytoplasm when compared to the nucleus (Fig 6F). GAPDH, HPRT1, and ACTB were found to be increased ∼5-fold in the cytoplasmic over the nuclear fraction. MKRN2 mRNA levels served as an additional control for our fractionation. As expected, we only observed a knockdown of MKRN2 mRNA levels in the cytoplasm, consistent with the subcellular localisation of the RISC complex. Upon siRNA knockdown of MKRN2, we observed a significant decrease in PA and NP cytoplasmic mRNA levels, while levels of GAPDH, HPRT1 and ACTB were unaffected (Fig 6F). These findings indicated to us that a loss of MKRN2 inhibits efficient export of viral mRNAs from the nucleus to the cytoplasm.

To confirm the persistence of this phenotype beyond the context of infection, a mini-replicon assay was performed in siRNA treated cells. Components of the influenza polymerase complex were transfected into HEK293T cells allowing the transcription of authentic viral mRNAs from a full length A/WSN/33 HA segment or a control mCherry mRNA pseudo segment carrying the HA packaging signal (psi) within the 5’ and 3’ UTRs. Consistent with previous results, upon fractionation we observed significantly less HA mRNA and mCherry-HApsi mRNA in the cytoplasmic fraction upon depletion of MKRN2 (Fig 6G). This indicated to us that MKRN2 association with viral mRNA is likely mediated by the 5’ and 3’ psi.

### The MKRN2-GLE1 complex contributes to IAV mRNA nuclear export

To decipher whether MKRN2 and GLE1 work in concert to mediate nuclear-cytoplasmic export of viral mRNA, we repeated the fractionation experiment in the context of GLE1 knockdown. Interestingly, knockdown of GLE1 phenocopied MKRN2 depletion, with both PA and NP mRNA levels significantly reduced in the cytoplasm, while again levels of GAPDH, HPRT1 and ACTB remained unchanged (Fig 6H). Additionally, GLE1 knockdown was confirmed by qPCR were a knockdown of expression occurred solely in the siGLE1 cytoplasmic fraction. To confirm that this observed reduction in cytoplasmic viral mRNA levels was reflected at the protein level, we performed a mini-replicon assay with the mCherry pseudo segment and measured the red fluorescence in cells depleted for MKRN2 or GLE1. Indeed, in the absence of MKRN2 or GLE1, we observed a significant reduction in mCherry signal. While GFP expressed from pcDNA 3.1 was unaltered under all conditions (Fig 6I). This demonstrates that reporter transgenes under a polII promoter are not affected by a loss of MKRN2.

Following on from this, we wanted to ascertain whether the disruption of nuclear-cytoplasmic trafficking due to a lack of MKRN2 was mediated by the coding sequence of IAV mRNAs or the packaging signal and conserved UTRs. In parallel, we aimed to identify whether transcription from the influenza viral polymerase was required to exploit MKRN2-mediated trafficking. To assess this, we designed PolII expression systems under a CMV promoter encoding for wild-type HA, NP or NA transgenes, or mScarlet and EBFP with the HA or NP packaging signals, respectively. We observed that both wild-type and reporter mRNAs demonstrated significantly reduced nuclear-cytoplasmic trafficking when MKRN2 was depleted (Fig 6J). Interestingly, for the wild-type CMV driven mRNAs we also observed a minor but significant increase in the nuclear fraction when MKRN2 expression was reduced. This led us to the conclusion that the 5’ and 3’ ends of IAV mRNA are necessary for MKRN2-associated nuclear export and that transcription via the viral polymerase is not required for MKRN2-mediated mRNA export.

## Discussion

Influenza A virus represents a highly infectious human pathogen ripe with potential for zoonotic spillover. IAV utilises a plethora of host proteins and pathways to successfully transcribe, translate and replicate its genome. A particularly fascinating aspect of IAV biology, in stark contrast to most other RNA viruses, is that RNA synthesis occurs in the nucleus. IAV mRNA synthesis is further complicated by the fact that, minimally, 8 out of the 10 predominant IAV mRNAs are not spliced.^20^ Splicing in eukaryotic cells generally leads to the direct recruitment of nuclear export factors such as NXF1 followed by the rapid trafficking of spliced mRNA to the cytoplasm.^21^ However, this is not the case for the majority of IAV transcripts, and successful replication necessitates other means to effectively export IAV mRNAs.

The export of IAV mRNAs, particularly the unspliced ones, is a poorly understood process. NXF1, the predominant mRNA export factor, has previously been linked with this process.^22^ A recent study has implicated a role for the TREX-2 complex in the nuclear-cytoplasmic trafficking of IAV mRNAs.^20^ The authors found that the nucleoporin, TPR, is important for this process via its recruitment of TREX-2 to the nuclear pore complex (NPC). For a subset of cellular mRNAs, and it seems IAV mRNAs, that are exported via this process, NXF1 is recruited by GANP, a component of TREX-2, which then leads to mRNA trafficking through the NPC.^20^ GLE1 is then involved in release of NXF1 which is then recycled back into the nucleus. An earlier study investigated the dependence of various known mRNA export factors for IAV mRNA export.^23^ The authors found that different IAV transcripts were either more or less reliant on cellular factors including NXF1 or UAP56. But beyond this work, little is known of RBPs that may be involved in the export of viral intronless mRNAs.

Techniques such as RIC, RAP-MS or VIR-CLASP represent a new possibility for a more comprehensive understanding of viral replication dynamics, including mRNA trafficking/export.^1,3,5^ Through a better understanding of the host proteome that is utilised and manipulated by viral proteins and nucleic acids, we may further understand not just important aspects of virus biology but also of cell biology more generally. We used the RIC technique to uncover RBPs that more readily associate to the mRNA pool within a cell during infection when compared to uninfected conditions. As the unspliced viral mRNA pool is a significant fraction of the total RNA pool,^11^ we reasoned that this technique would be an ideal approach to identify a novel host factor involved in the nuclear export of IAV mRNAs. Using 2 different IAV strains, we hoped to reduce the background and uncover a cellular protein whose role is conserved across IAV strains.

In our initial analysis of our MS datasets, we confirmed the substantial enrichment of the ubiquitous and highly expressed viral protein, NP, in both datasets (Fig 2A & 2B), as expected. We next looked to validate some of the more interesting RBPs that we found to be enriched on mRNAs from IAV infected cells. Here, our aim was to determine which, if any, played a crucial role in viral replication. Using a fluorescent reporter virus, we found that MKRN2 depletion appeared to quash viral protein expression, while a similar protein from the same family, MKRN1, was not found to be required for viral protein expression. However, previous data surrounding the emergence of the *MKRN2* gene make this result less surprising than at first thought. *MKRN2* appears to have arisen from a gene duplication of *MKRN1* over 450 million years ago.^24,25^ Phylogenetic analysis has shown that for most of its history, the *MKRN2* gene lacks any significant evolutionary divergence. In fact, when the MKRN1 and MKRN2 proteins of human, mouse, zebrafish and yellowfin tuna were analysed for sequence conservation, each was found to cluster with its own orthologues, forming 2 distinct paralogous groups.^24^ This implies that MKRN1 and MKRN2 likely have distinct roles within the cell, while the lack of evolutionary divergence between orthologues hints that MKRN2 has a conserved cellular role among vertebrates.

In 2020 Wolf et al. published one of the only mechanistic studies to date into the potential role of MKRN2 as an RNA-binding protein. Their work indicated that MKRN2 binds to a subset of cellular mRNAs, but that in general MKRN2 acts to halt nuclear export of these mRNAs to the cytoplasm. The authors also identified GLE1 as an interacting partner of MKRN2. They speculate that binding of MKRN2 to an mRNA blocks GLE1 association with said mRNA and prevents efficient nuclear export.^15^ We performed our own analysis of the same IP-MS raw data to uncover any novel interactors that are unique to MKRN2, over MKRN1 and MKRN3 that may have been overlooked in the initial study (Fig 6A). As we have already shown that a loss of MKRN1 does not phenocopy a loss of MKRN2 in the context of IAV replication, we reasoned that identifying MKRN2-specific interactors would aid in developing more mechanistic insights. On analysis, we found GLE1 alongside several other proteins described by Wolf et al., that are directly involved in RNA transport and nuclear export; ZC3H14, RAE1, FYTTD1 and NUP42 (Fig 6B & C). In addition though, we uncovered 2 components of the exon junction complex, MAGOH and SAP18, that associate with MKRN2 but not MKRN1 or MKRN3. MAGOH is an RBP and has been reported to bind to the known mRNA export factor NXF1. Interestingly, MAGOH remains associated with bound mRNAs even after nuclear export.^26^ While SAP18 has previously been described as a late associating factor of the exon junction complex.^27^ Taken together, this list of interactors is compelling evidence that MKRN2 is involved in nuclear export of bound RNA, but that MKRN2 would likely positively regulate nuclear export as it distinctly associates with many well-established components of mRNA nuclear export. This is further validated herein, where we find a significant cytoplasmic depletion phenotype for IAV transcripts when expression of either MKRN2 or its co-factor GLE1 is reduced in infected cells.

As stated above, it has already been shown that GLE1 is involved in the recycling of NXF1 back into the nucleus,^21^ and that MKRN2 has been validated previously as a GLE1 interacting protein.^15^ With recent work demonstrating a role for TREX-2 mediated recruitment of NXF1 to viral mRNAs for efficient nuclear export,^20^ it may be important to investigate any potential links between MKRN2 and TREX-2, in the presence and absence of virus infection. In fact, the IP-MS published by Wolf et al (2020) demonstrates that MKRN2 associates with FYTTD1, a UAP-56 interacting protein, a known interactor of the TREX complex. We have already demonstrated herein that a reduction in FYTTD1 reduced IAV infection kinetics (Fig 3B). Taken together, these data may implicate MKRN2 as an RNA-binding mRNA export adapter protein for a subset of cellular and viral mRNAs. Previously, two splicing factors 9G8 and SRp20 have been described as export adapter proteins, through binding to target mRNA and direct recruitment of NXF1 to mediate mRNA export.^28^ Therefore, it is possible that MKRN2 may be important for the recruitment of the export complex, explaining how a loss of MKRN2 prevents efficient nuclear export of IAV mRNAs.

We have shown that MKRN2-mediated viral mRNA export is indeed occurring, at least for the primary transcripts generated upon IAV infection. It is likely though that this binding and nuclear export continues throughout infection as MKRN2 association appears to be mediated by the sequence of the transcript itself. With all this in mind, we are proposing a model for IAV mRNA nuclear export. Upon IAV infection MKRN2 protein begins to accumulate in the nucleus of infected cells. This nuclear MKRN2 then binds to IAV transcripts and potentially recruits export adapter proteins such as FYTTD1 of MAGOH. This in turn recruits export machinery such as TREX leading to trafficking to the NPC, where the complex interacts with GLE1, and viral mRNAs are efficiently exported to the cytoplasm. What is clear from the work presented here is that MKRN2 is an important RBP in the export of IAV mRNAs from the nucleus and that MKRN2 likely carries out this role through its proven association with members of the mRNA export machinery, of which GLE1 is one.

**Figure 7 –.**
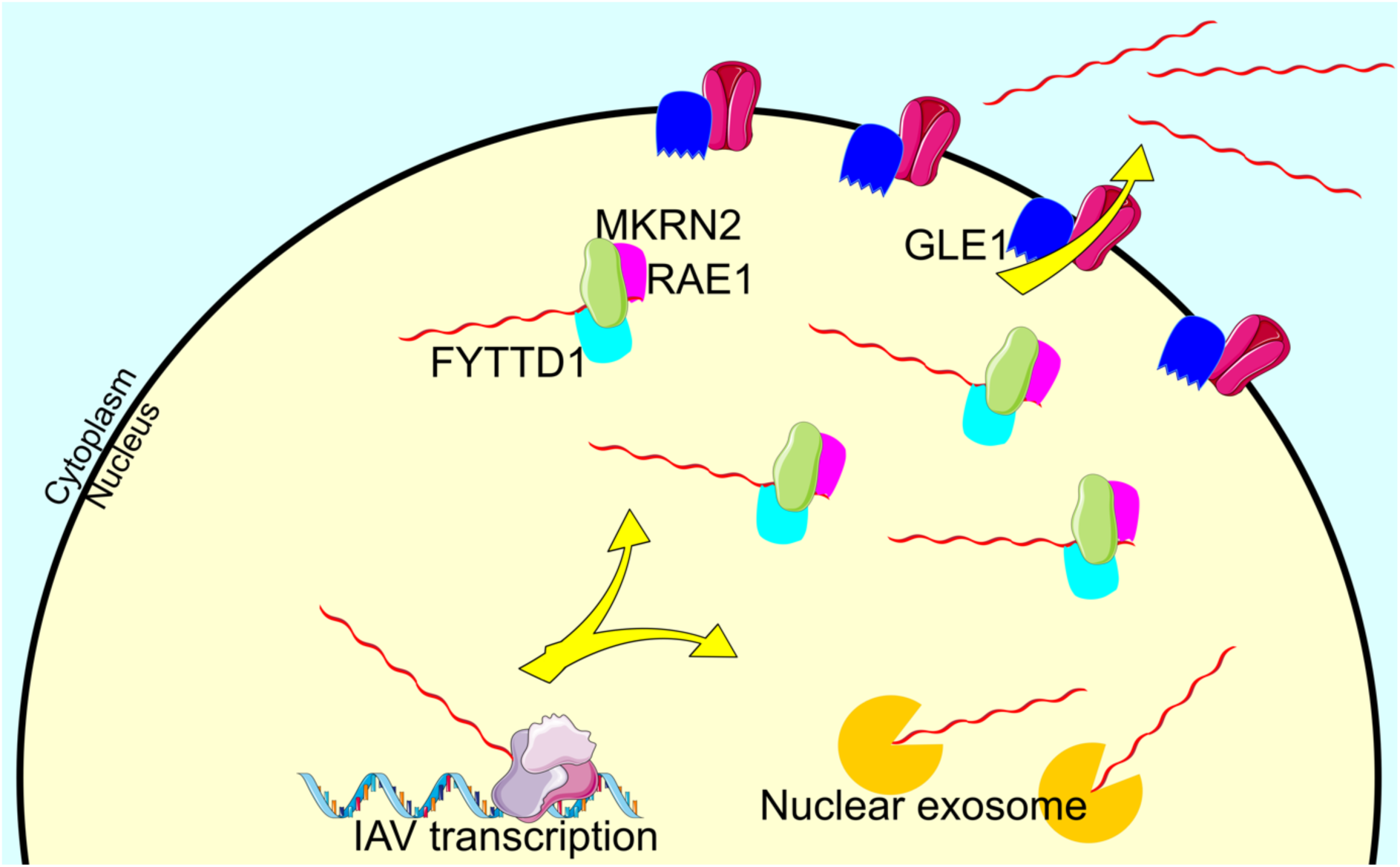
Model of MKRN2 mediated nucleo-cytoplasmic shuttling of IAV mRNA. MKRN2 likely associates with GLE1 and potentially several other cellular components normally involved in mRNA nuclear export such as RAE1 and FYTTD1. Without this association IAV transcripts remain confined to the nucleus, which likely inevitably leads to degradation by the nuclear exosome.

## Methods

### Cells

For this work immortalised A549 (ATCC; CCL-185), HEK 293T (ATCC; CRL-3216) and MDCK (CCL-34) cells were used. All cells were cultured in DMEM supplemented with 1% Pen-Strep (Thermo Fisher Scientific; 15140122) and 5% FBS (Thermo Fisher Scientific; 10270106) at 37°C and 5% CO_2_.

### Virus stocks and infections

WSN (A/WSN/33) viral stocks were generated from a reverse genetics system that has been described previously.^29^ WSN-PA_mNeon virus was generated from the same reverse genetics system, where the pPolI-PA plasmid was substituted for the pPolI-PA_mNeon plasmid encoding for a C-terminal mNeon followed by a duplicated packaging signal. Cal09 (A/California/7/2009) and Norway14 (A/Norway/466/2014) viruses were grown from isolates acquired from the National Institute for Biological Standards and Control, UK. All stocks were grown on MDCK cells in IAV growth media consisting of DMEM, Pen-Strep, 0.2% BSA (Merck; A8412), 25mM HEPES (Merck; H0887) and TPCK-trypsin (Merck; T8802), and titred on MDCK cells by plaque assay.

### Influenza RIC

The protocol utilised here for the capture of the RNA interactome of poly(A)+ RNA (RIC) has been described previously.^1,7^ Herein, we use all the same lysis and wash buffers and timepoints for the harvesting of poly(A)+ RNA, and by extension, the RBPs associated with them. In short, A549 cells were seeded in 15cm plates prior to infection. Cells were either mock infected or infected with A/WSN/33 at an MOI 3 PFU/cell, or infected with A/California/7/2009 at an MOI 10 PFU/cell for 1 h in DMEM. Infected cells were incubated in IAV growth media for 16 h before undergoing UV crosslinking at 254nm and lysis. Oligo dT beads (NEB; S1419S) were used to pull down poly(A)+ mRNAs and covalently bound RBPs. After a series of wash steps described in the published protocol, a total of 10% of the pulldown was kept for RNA isolation, with RNA being eluted by boiling in dH_2_O followed by a proteinase K digestion (Thermo Fisher Scientific; 25530049). RBPs were eluted for Western blotting, silver staining (Thermo Fisher Scientific; 24612) and mass spectrometry by an RNase digest. Protein isolates for mass spectrometry were precipitated, reduced, trypsin digested followed by TMT labelling (Thermo Fisher Scientific; 90111), following the manufacturer’s protocol.

### Western blot

Western blotting was performed as has been previously described.^30^ Primary antibodies for influenza NP (Abcam; PA5-32242), PA (Thermo Fisher Scientific; Pa5-32223), NS1 (Santa Cruz; Sc-13056), YTHDF2 (Proteintech; 24744-1-AP), ACTB (Proteintech; 66009-1-ig), Flag (Proteintech; 66008-3-ig) and MKRN2 (Novus Biologics; nbp2-17301) were all used at a 1:5000 dilution. Proteins bands on blots were visualised by chemiluminescence.

### LC-MS/MS and data analysis

Mass spectrometry of RIC samples was performed at the Cambridge Centre for Proteomics. LC-MS/MS analyses were performed on an Orbitrap™ Fusion™ Lumos™ Tribrid™ mass spectrometer and performed as previously described. Raw mass spectrometry data was analysed using MaxQuant software (v2.4.7.0) as has been previously described.^31^ Each individual replicate was aligned to the human Uniprot database^32^ and the corresponding influenza virus Uniprot protein database. Statistical analysis was performed using Perseus software (v2.0.11) as previously described, and data was normalised using an internal reference. Potential contaminants were removed, proteins had to be present in at least 2 replicates within a single group to be included, a t-test was used to calculate significance with p<0.05.

### siRNA knockdowns

For all siRNA knockdowns, a pair of siRNAs targeting the specified mRNA, or a non-specific control siRNA were used. These siRNA sequences are listed in Supplementary Table 2. A final concentration of 20nM total siRNA was used in each condition, and siRNAs were all transfected using Lipofectamine™ RNAiMAX (Invitrogen; 13778075) following the manufacturer’s instructions. A549 or 293T cells were seeded at the time of transfection and incubated for 24 h before media was changed for fresh growth media. Cells were then incubated for a further 24 h prior to either infection of transfection.

### CLIP-qPCR

The CLIP-qPCR protocol employed here has been published previously.^33^ Briefly, 293T cells were seeded one day prior to transfection. Cells were then transfected with a pCI plasmid encoding for either 3x Flag tagged mCherry or MKRN2, with mCherry acting as a negative control. At 48 h post transfection cells were infected with A/WSN/33 at an MOI 3 PFU/cell and incubated for 8 h. At 8 hpi media was removed from the cells, before being washed in ice cold PBS. Cells then underwent UV crosslinking at 254nm. Cells were then scraped, pelleted and lysed in 250 µl of NP40 lysis buffer (HEPES-KOH pH 7.5 50mM, KCl 150mM, EDTA pH 8.0 2mM, NaF 1mM and NP40 0.5%) on ice for 30 min. Lysates underwent clarification by centrifugation and 10% was then removed as an input and boiled in Laemmli buffer at 95°C for 10 min for western blotting. The remaining 90% of lysate underwent immunoprecipitation with anti-Flag antibody (Merck, F1804) coupled to Protein G Dynabeads (ThermoFisher, 10004D) on a rotator for 1 h at room temperature. Beads were then captured, washed five times with 1 mL NP40 lysis buffer and three times with PBS. A fraction (10%) of the beads was removed, resuspended in Laemmli buffer and boiled at 95°C for 10 min for Western blotting. The remaining 90% of beads underwent proteinase K digestion (ThermoFisher, EO0491) at 55°C for 60 min to release bound RNA, following the manufacturer’s instructions. RNA was then isolated from the digestion by TRIzol LS (ThermoFisher, 10296028) and ethanol precipitation. Isolated RNA underwent cDNA conversion using oligo dT or A/WSN/33 specific primers^34^ (Supplementary Table 1). Quantification of the cDNAs was performed by qPCR, described below. Fold enrichment of RNA levels was quantified using the ΔΔCT method and normalised to input mCherry levels.

### Establishment of a MKRN2 overexpression cell line using a lentiviral vector

A lentiviral expression plasmid was generated encoding for full length wild-type MKRN2 C-terminally tagged with 3 Flag epitope tags. MKRN2 was followed by a P2A self-cleaving peptide and a puromycin resistance gene, similar to a system we have published previously.^35^ Again, similar to previous publications, we rescued lentiviral particles by co-transfecting 293T cells with the MKRN2 lentiviral vector alongside d8.74 (Addgene; 22036) and MD2G (Addgene; 12259). At 72 h post-transfection the supernatant was harvested, passed through a 0.45 μm filter and then overlaid onto fresh A549 cells. At 72 h post transduction cells were treated with 2 μg/ml puromycin and left to select for 72 h. Cells were then single cell cloned in 96 well plates in the presence of 2 μg/ml puromycin and left to grow for 3 weeks. Single cell colonies were then selected, expanded, and checked by Western blot to confirm the expression of the MKRN2 transgene.

### IAV multicycle titres

For multicycle IAV infections, A549 cells were seeded and transfected with siRNA simultaneously, as described above. At 48 h post transfection cells were infected with either A/WSN/33 at an MOI 0.01 PFU/cell, or A/California/7/2009 or A/Norway/466/2014 at an MOI 0.1 PFU/cell in DMEM for 1 h. The inoculum was then removed, and cells incubated for 48 h for A/WSN/33 or 72 h for A/California/7/2009 and A/Norway/466/2014 in IAV growth media, described above. Supernatants were then titred for influenza virus on MDCK cells by plaque assay.

### IAV qPCR growth curves

A549 cells were infected with A/WSN/33 or A/California/7/2009 virus at an MOI 3 of 10 PFU/cell, respectively. At 0, 2, 4 and 6 hpi RNA was extracted from cells using TRIzol (Invitrogen; 15596026). RNA was isolated and precipitated following the manufacturer’s institutions and 200ng was reverse transcribed into cDNA using the ABI cDNA synthesis kit (Applied Biosystems; 4368814). For all cellular targets RT was performed using an oligo dT, while viral RNA underwent separate RT reactions with primers specific to each mRNA or vRNA.^34^ These primers are listed in Supplementary Table 1 alongside the primers used for qPCR amplification. All qPCR experiments were performed using SYBR Select Master Mix (Applied Biosystems; 4472908) following the manufacturer’s instructions. All qPCR data were quantified using the ΔΔCT method.

### Nucleo-cytoplasmic fractionation

A nuclear-cytoplasmic fractionation was used to determine the differences in viral and cellular mRNA abundance between these 2 cellular compartments. Each time this was performed in an infected context cells were treated with 100 μg/ml cycloheximide at 1 hpi so that only primary transcription was observed and quantified. We have published this method previously for virus infected cells,^33^ but in short, cells that had been previously treated with siRNA and infected were washed in ice cold PBS at the desired timepoint post-infection. Cells were scraped from the plate and pelleted at 4°C. PBS was removed, and cells were resuspended in NP40 lysis buffer (10mM Tris-HCl pH 7.4, 10mM NaCl, 3mM MgCl2 and 0.5% NP-40), followed by brief vortexing and incubation on ice for 5 minutes. Lysates were centrifuged at max speed (14,000g) for 10 seconds to pellet the nuclei. This supernatant was removed and labelled as the cytoplasmic fraction. The nuclei pellet was washed in a further volume of NP-40 lysis buffer before being pelleted again by centrifugation for 10 seconds. The supernatant was discarded, and the pellet was resuspended in either Laemmli buffer or TRIzol depending on whether protein or RNA was required. RNA samples then underwent reverse transcription followed by qPCR to quantify viral or cellular transcript levels. These data were quantified using the ΔΔCT method and normalised to the levels of each mRNA species in the nuclear fraction of control siRNA treated cells.

### Immunofluorescence and analysis

A549 cells were seeded onto coverslips 24 h prior to infection. Cells were infected with A/WSN/33 at MOI 1, and the infection was synchronised by incubation at 4°C for 1 h. Mock, 4 and 8 hpi samples were fixed with PFA and processed for immunofluorescence. In brief, cells were fixed in 4% PFA, washed in PBS and permeabilised with 0.2% Triton X-100 for 10 minutes. Cells were blocked in 3% BSA prior to incubation with primary antibody dilutions for 2 h. A 1:500 dilution of NP antibody (Abcam; PA5-32242) and a 1:200 dilution of MKRN2 antibody (Novus Biologics; nbp2-17301). After PBS washes the cells were incubated in secondary antibody for 1 h. Anti-mouse secondary was used at 1:500 (Thermo Fisher Scientific; A28175) and anti-rabbit secondary was used at 1:500 (Thermo Fisher Scientific; A-21245). Finally cells were counterstained with Hoechst and mounted prior to imaging.

Coverslips were imaged with a Leica Stellaris 8 confocal microscope with a 63x objective with 1.4 NA, resulting in a pixel size of 180nm. Single slices of entire fields of view were acquired for more than 10 cells per condition and analysed with an automated Fiji macro. Briefly, after background subtraction, nuclei and cytoplasm were thresholded and annotated based on the Hoechst and Phalloidin signals respectively. Then, integrated fluorescence intensity values for each were measured and ratios for each single cell were calculated. In total 25, 11 and 33 cells were assessed by this method for Mock, 4 and 8 hpi samples, respectively, from a single coverslip. Statistical significance between conditions was calculated by one-way ANOVA.

### IAV mini-replicon assay

293T cells were first transfected with siRNA and incubated for 72 h. Then they were transfected with equal amounts of pcDNA-PB2, pcDNA-PB1, pcDNA-PA and pcDNA-NP and 1 reporter expressing plasmid either, pPolI-HA, pPolI-mCherry_HApsi. Cells being used for fluorescence imaging and quantification were also transfected with a pcDNA-GFP plasmid, where the GFP was used as a transfection control. Transfected cells were then incubated for 24h before either imaging for number of fluorescent cells, or nuclear cytoplasmic fractionation followed by RNA extraction and qPCR.

### Statistical analysis

Unless otherwise stated, all experiments were performed as 3 independent biological replicates. When bar graphs are used to represent the data, each individual replicate can be observed as a point overlaid on the bar. Significance was calculated by performing Student’s t-test with * indicating p<0.05, ** indicating p<0.01 and *** indicating p<0.001.

## Acknowledgements

This research was funded in part by an ERC-STG grant, PTFLU 949506 awarded to D.G.C. This research received infrastructure support from the Wellcome-Wolfson Institute for Experimental Medicine at Queen’s University Belfast.

## Supplemental figures

**Supplementary Figure 1.**
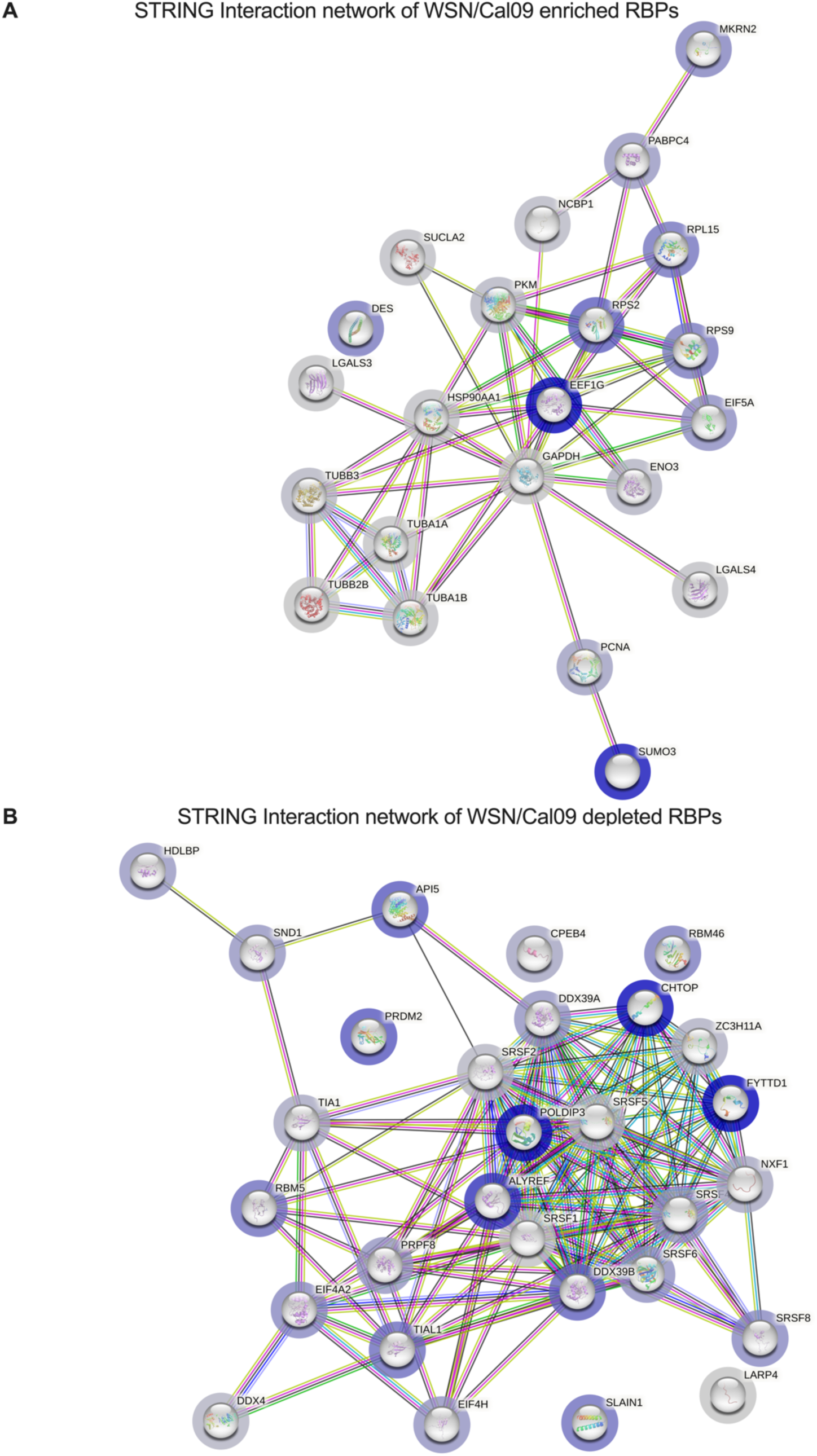

**Supplementary Figure 2.**
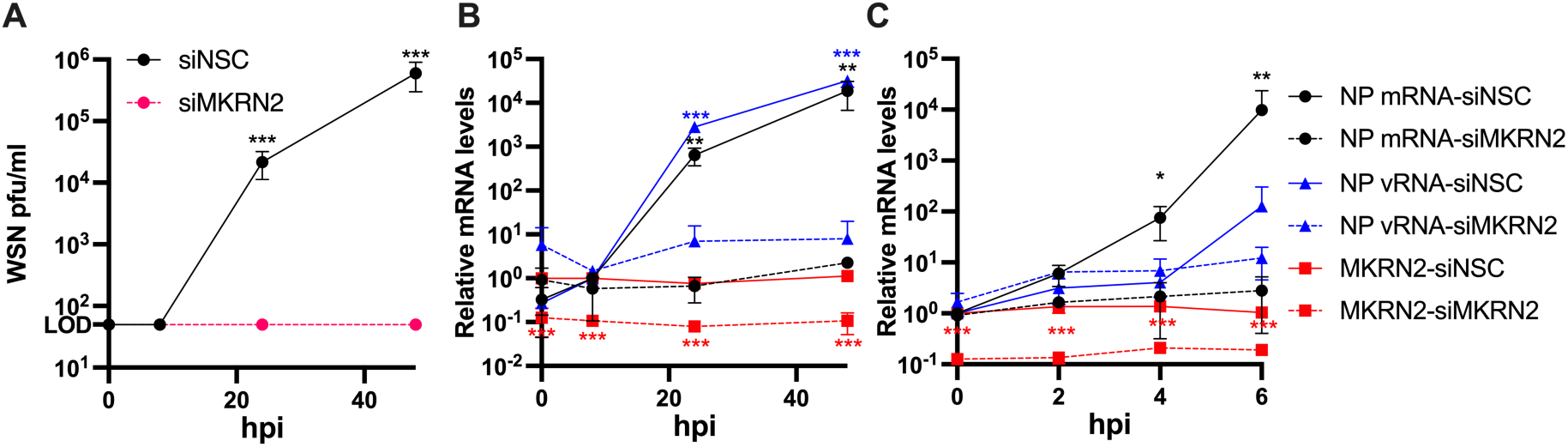

**Supplementary Figure 3.**
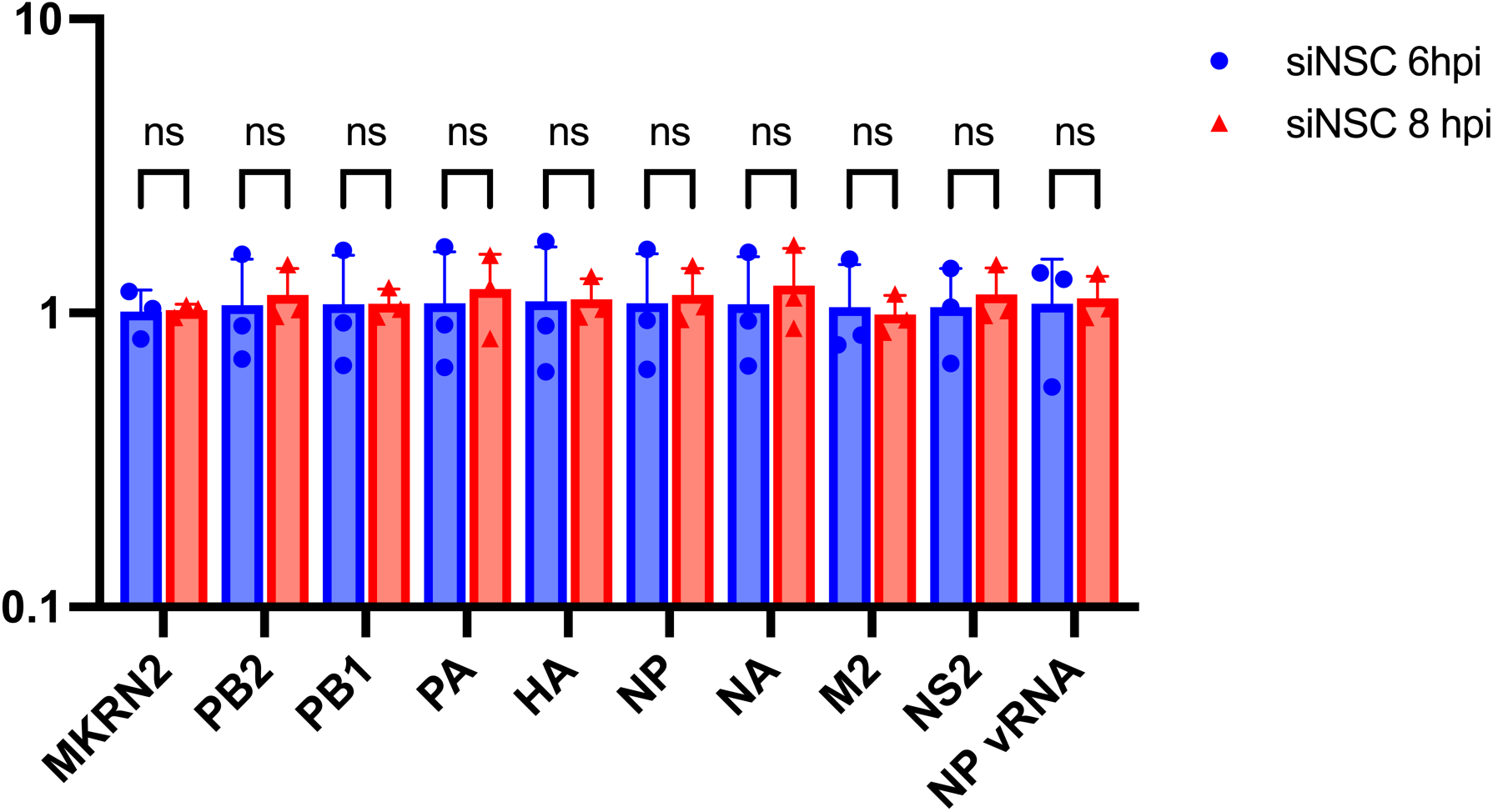

